# Human cortical slices reveal pulse-number-dependent synaptic plasticity during theta-burst magnetic stimulation

**DOI:** 10.64898/2026.10.23.761311

**Authors:** Christos Galanis, Jakob Straehle, Zsolt Turi, Jürgen Beck, Andreas Vlachos

## Abstract

Theta-burst transcranial magnetic stimulation is widely used to modulate human brain activity in clinical and experimental settings, yet how clinically relevant stimulation protocols engage synaptic plasticity in human cortical circuits remains poorly understood. Here, we combined controlled magnetic stimulation of viable adult human cortical slices with single-cell electrophysiology to determine whether pulse number constrains synaptic plasticity during intermittent theta-burst stimulation (iTBS). As a functional reference, 10 Hz repetitive magnetic stimulation delivered with 900 pulses robustly potentiated excitatory synaptic transmission in human cortical pyramidal neurons. Similarly, iTBS900 significantly potentiated AMPA receptor-mediated miniature excitatory postsynaptic currents. In contrast, the clinically established 600-pulse iTBS protocol failed to reliably induce detectable synaptic plasticity under otherwise identical stimulation conditions. Paired within-patient experiments confirmed distinct synaptic responses to iTBS600 and iTBS900 in the same human cortical tissue preparations. Together, these findings identify pulse number as an important determinant of theta-burst magnetic stimulation-induced synaptic plasticity in human cortical microcircuits and highlight stimulation dose as a relevant parameter for the design and refinement of therapeutic stimulation protocols.

## Introduction

Theta-burst transcranial magnetic stimulation is widely used to modulate human brain activity in clinical and experimental settings (Massimini et al., 2026; Siebner et al., 2022). Theta-burst stimulation, and in particular intermittent theta-burst stimulation (iTBS), has become an established stimulation paradigm because it induces lasting physiological effects while substantially reducing stimulation time compared to conventional 10 Hz repetitive transcranial magnetic stimulation protocols (Huang et al., 2005; Huang & Rothwell, 2004). Despite its widespread use, how clinically relevant theta-burst stimulation protocols engage synaptic plasticity in human cortical circuits remains poorly understood.

Experimental studies in rodents and organotypic brain tissue cultures have demonstrated that repetitive magnetic stimulation induces long-lasting functional and structural plasticity at excitatory and inhibitory synapses (Galanis et al., 2024, 2025; Lenz et al., 2015, 2016; Vlachos et al., 2012). Recent computational and experimental work further suggests that stimulation-induced plasticity depends critically on stimulation dose and neuronal excitability state (Anil et al., 2023; Bai et al., 2023; Baker et al., 1995; Hananeia et al., 2025; Momi et al., 2022; Zrenner et al., 2018). Among these parameters, pulse number has emerged as an important determinant of synaptic plasticity induced by repetitive magnetic stimulation (Galanis et al., 2026). Whether pulse-number similarly shapes synaptic responses to clinically relevant theta-burst stimulation in viable adult human cortical circuits remains unknown.

To address this question, we combined controlled magnetic stimulation of acute adult human cortical slices with single-cell electrophysiological recordings. Using clinically relevant theta-burst stimulation paradigms under otherwise identical stimulation conditions, we tested whether total pulse number influences the induction of synaptic plasticity in human cortical pyramidal neurons.

## Results

### Repetitive magnetic stimulation induces excitatory synaptic potentiation in human cortical slices

To establish whether repetitive magnetic stimulation induces synaptic plasticity in viable adult human cortical circuits, acute cortical slices prepared from surgical tissue resections were stimulated under controlled ex vivo conditions (Figure 1). Slices were positioned directly above a 70-mm inner-diameter figure-of-eight magnetic stimulation coil, with stimulation applied from below (Figure 1A). Finite element modeling was used to estimate the induced electric field strength within the tissue under the stimulation configuration (90.5 V/m). Sister slices from the same patient tissue preparations served as matched sham controls throughout all experiments.

**Figure 1:**
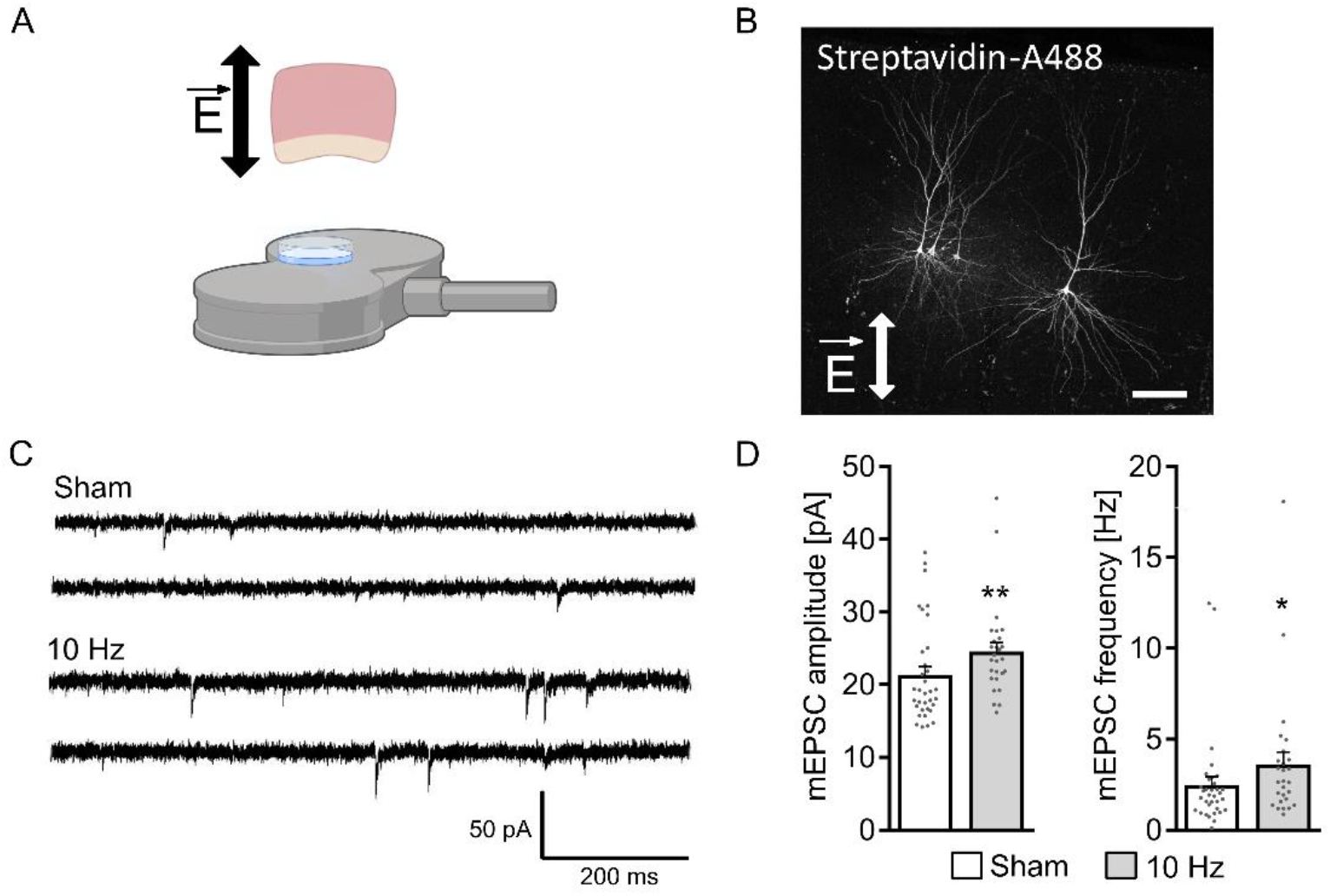
10 Hz repetitive magnetic stimulation induces excitatory synaptic potentiation in human neocortical pyramidal neurons. (A) Schematic illustration of a human neocortical slice showing the direction of the induced electric field (top) and the figure-of-eight coil positioned below the Petri dish during stimulation (bottom). (B) Representative biocytin-filled layer II/III pyramidal neurons visualized with Alexa Fluor 488-conjugated streptavidin. Scale bar, 200 μm. (C, D) Sample traces and group data of AMPA receptor-mediated miniature excitatory postsynaptic currents (mEPSCs) recorded from layer II/III pyramidal neurons in sham and 10 Hz stimulated slices 2-4 h after stimulation. mEPSC amplitude: sham, 21.4 ± 1.2 pA (n = 34 neurons from 4 patients); 10 Hz, 24.6 ± 1.2 pA (n = 27 neurons from 4 patients); Mann-Whitney U = 280. mEPSC frequency: sham, 2.5 ± 0.5 Hz (n = 34 neurons from 4 patients), 10 Hz: 3.6 ± 0.7 pA (n = 27 neurons from 4 patients); Mann-Whitney U = 302. Data are mean ± SEM. *p < 0.05; **p < 0.01.

As a functional reference condition, we first applied a previously established 10 Hz repetitive magnetic stimulation protocol consisting of 900 pulses, which induces synaptic plasticity in mouse hippocampal tissue cultures (Galanis et al., 2024, 2025; Lenz et al., 2015; Vlachos et al., 2012). Whole-cell voltage-clamp recordings were obtained from layer II/III pyramidal neurons 2–4 h after stimulation (Figure 1C–D). AMPA receptor-mediated miniature excitatory postsynaptic current (mEPSC) amplitude and frequency were significantly increased in stimulated compared with patient-matched sham slices (Figure 1D). These findings establish that viable adult human cortical circuits express repetitive magnetic stimulation-induced synaptic potentiation under the present experimental conditions.

### iTBS900 induces excitatory synaptic plasticity in human cortical pyramidal neurons

After establishing that repetitive magnetic stimulation induces synaptic potentiation in human cortical slices under the present experimental conditions, we next examined whether intermittent theta-burst stimulation (iTBS) engages excitatory synaptic plasticity in human cortical microcircuits (Figure 2). Human cortical slices were stimulated using an iTBS protocol consisting of 900 pulses (iTBS900) while maintaining the same stimulation geometry and estimated electric field strength as in the 10 Hz reference condition (Figure 2A).

**Figure 2:**
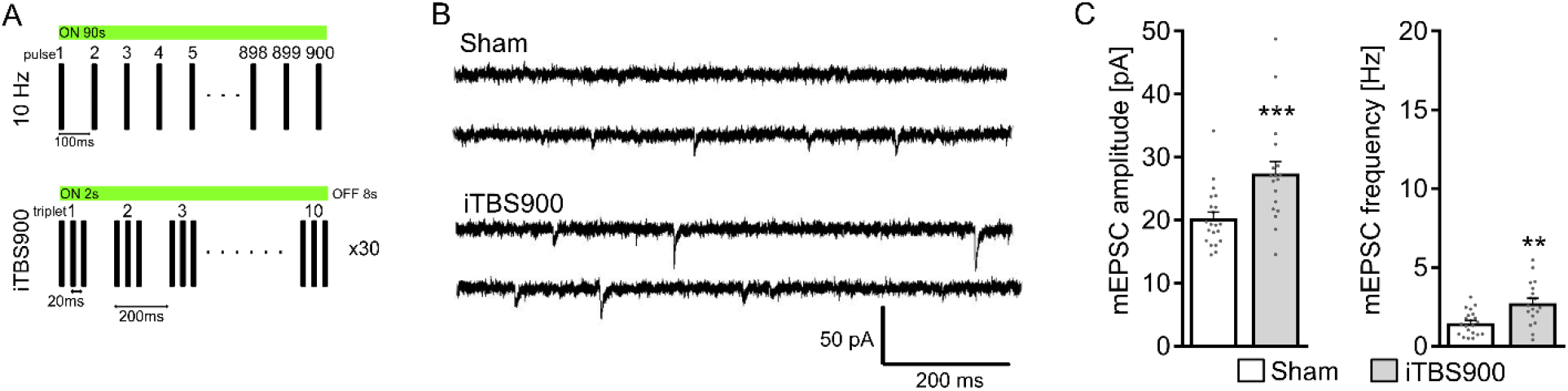
iTBS900 induces excitatory synaptic potentiation in human neocortical pyramidal neurons. (A) Schematic illustration of the 10 Hz and iTBS900 protocols. (B, C) Sample traces and group data of AMPA receptor-mediated miniature excitatory postsynaptic currents (mEPSCs) recorded from layer II/III pyramidal neurons in sham and iTBS900-stimulated slices 2-4 h after stimulation. mEPSC amplitude: sham, 20.3 ± 1.0 pA (n =21 neurons from 4 patients); iTBS900, 27.4 ± 1.9 pA (n = 18 neurons from 4 patients); Mann-Whitney U = 73. mEPSC frequency: sham, 1.5 ± 0.2 Hz (n =21 neurons from 4 patients); iTBS900, 2.8 ± 0.3 Hz (n = 18 neurons from 4 patients); Mann-Whitney U = 84. Data are mean ± SEM. **p < 0.01; ***p < 0.001.

Whole-cell voltage-clamp recordings obtained from layer II/III pyramidal neurons 2–4 h after stimulation revealed a significant potentiation of AMPA receptor-mediated mEPSCs in iTBS900-stimulated slices compared with patient-matched sham slices (Figure 2B, C). Both mEPSC amplitude and frequency were significantly increased following iTBS900, demonstrating potentiation of excitatory synapses in viable human cortical circuits.

### iTBS600 and iTBS900 produce distinct synaptic responses in human cortical slices

We next asked whether the clinically established 600-pulse iTBS protocol induces comparable synaptic plasticity in viable human cortical circuits under otherwise identical stimulation conditions (Figure 3A). In contrast to iTBS900, whole-cell recordings obtained from layer II/III pyramidal neurons 2–4 h after stimulation revealed no significant change in AMPA receptor-mediated mEPSC amplitude or frequency compared with patient-matched sham slices (Figure 3B, C). Thus, under otherwise identical stimulation conditions, iTBS600 did not reliably produce detectable excitatory synaptic potentiation.

**Figure 3:**
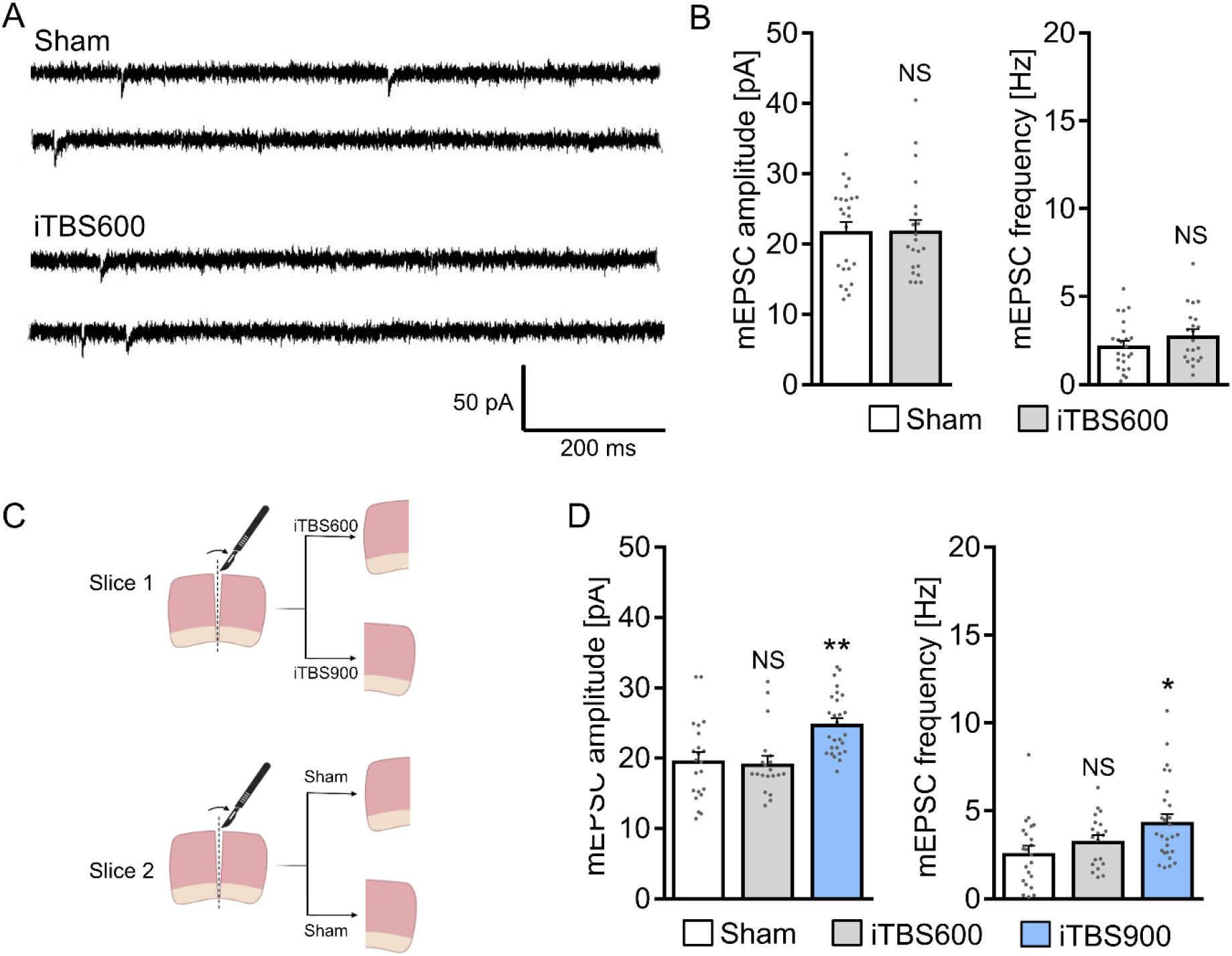
iTBS600 and iTBS900 produce distinct synaptic responses inhuman neocortical pyramidal neurons. (A, B) Sample traces and group data of AMPA receptor-mediated miniature excitatory postsynaptic currents (mEPSCs) recorded from layer II/III pyramidal neurons in sham and iTBS600-stimulated slices 2-4 h after stimulation. mEPSC amplitude: sham, 21.9 ± 1.3 pA (n =24 neurons from 4 patients); iTBS600: 21.9 ± 1.5 pA (n = 21 neurons from 4 patients); Mann-Whitney U = 240. mEPSC frequency: sham, 2.2 ± 0.3 Hz (n =24 neurons from 4 patients); iTBS600: 2.8 ± 0.3 pA (n = 21 neurons from 4 patients); Mann-Whitney U = 198. (C) Schematic illustration of the within-patient experimental design. Neocortical slices obtained from the same patient tissue preparations were assigned to sham, iTBS600, or iTBS900 condition, enabling direct comparison of the stimulation conditions within individual patients. (D) Group data of AMPA receptor-mediated miniature excitatory postsynaptic currents (mEPSCs) recorded from layer II/III pyramidal neurons in patient-matched sham, iTBS600, and iTBS900 slices 2-4 h after stimulation. mEPSC amplitude: sham, 19.6 ± 1.2 pA (n =22 neurons from 3 patients); iTBS600, 19.2 ± 1.0 pA (n = 20 neurons from 3 patients); iTBS900, 24.9 ± 0.8 pA (n = 27 neurons from 3 patients). mEPSC frequency: sham, 2.6 ± 0.4 Hz (n =22 neurons from 3 patients); iTBS600, 3.3 ± 0.3 Hz (n = 20 neurons from 3 patients); iTBS900, 4.4 ± 0.4 pA (n = 27 neurons from 3 patients); Kruskal-Wallis test. Data are mean ± SEM. NS, not significant; *p < 0.05; **p < 0.01.

To directly assess the influence of pulse number while minimizing interpatient-variability, we compared iTBS600 and iTBS900 in matched tissue preparations obtained from the same patients (Figure 3D). In these within-patient experiments, iTBS900 consistently induced potentiation of AMPA receptor-mediated mEPSCs relative to patient-matched sham slices, whereas iTBS600 did not produce detectable synaptic potentiation in the same cortical tissue preparations. Together, these findings demonstrate distinct synaptic responses to iTBS600 and iTBS900 and identify pulse number as an important determinant of theta-burst magnetic stimulation-induced synaptic plasticity in viable adult human cortical circuits.

## Discussion

In the present study, we combined controlled magnetic stimulation of viable adult human cortical slices with single-cell electrophysiology to examine how pulse number influences the induction of synaptic plasticity during clinically relevant theta-burst stimulation. We demonstrate that iTBS with 900 pulses induce excitatory synaptic potentiation in adult human cortical microcircuits, whereas the clinically established iTBS protocol with 600 pulses did not reliably produce detectable synaptic potentiation under otherwise identical stimulation conditions. Importantly, within-patient experiments revealed distinct synaptic responses to iTBS600 and iTBS900 in tissue preparations obtained from the same patient. Together these findings identify pulse number as an important determinant of theta-burst magnetic stimulation-induced synaptic plasticity in human cortical circuits.

Previous experimental work in rodents and organotypic tissue cultures has demonstrated that repetitive magnetic stimulation induces functional and structural synaptic plasticity through activity-dependent mechanisms involving NMDA receptors, voltage-gated calcium channels, and coordinated pre- and postsynaptic signaling (Galanis et al., 2025; Lenz et al., 2015, 2016; Vlachos et al., 2012). Building on these findings, recent experimental and computational studies suggest that stimulation-induced plasticity is sensitive to cumulative stimulation dose and may exhibit threshold-like behavior depending on neuronal excitability state and stimulation intensity (Galanis et al., 2025; Hananeia et al., 2025). Our findings extend these observations to viable adult human cortical circuits by demonstrating that synaptic responses to theta-burst magnetic stimulation depend on pulse number. Both conventional 10 Hz stimulation delivered with 900 pulses and iTBS900 potentiated excitatory synaptic transmission, whereas iTBS600 did not reliably produce detectable synaptic potentiation under otherwise identical stimulation conditions. These observations are consistent the possibility that cumulative pulse number contributes to threshold-like engagement of synaptic plasticity in human cortical circuits.

An important implication of these findings is that stimulation dose may contribute to variability in the engagement of synaptic plasticity by clinically applied iTBS protocols. Considerable interindividual variability in physiological and clinical responses to iTBS has been reported in human studies, yet the biological basis of this variability remains incompletely understood. Our data suggest that pulse number represents one parameter that can influence whether detectable synaptic potentiation is induced in human cortical circuits. In this context, accelerated neuromodulation approaches such as the Stanford Accelerated Intelligent Neuromodulation Therapy (SAINT) are of particular interest, as they combine a higher pulse number per session with repeated daily iTBS sessions and individualized targeting (Cole et al., 2020). While the present experiments do not model repeated clinical stimulation or establish a relationship between pulse number and therapeutic efficacy, they provide a mechanistic framework for considering stimulation dose as one factor that may influence engagement of plasticity-related processes in human cortical networks.

Importantly, the present findings also demonstrate that viable adult human cortical tissue can be interrogated using controlled magnetic stimulation paradigms combined with direct single-cell electrophysiological recordings. Human cortical slice models provide an opportunity to bridge mechanistic experimental neuroscience and human neuromodulation research by enabling direct investigation of stimulation-induced synaptic plasticity in native human cortical microcircuits under controlled conditions. In contrast to indirect physiological readouts commonly used in vivo, this approach permits direct assessment of synaptic function at cellular resolution while preserving key features of human cortical circuit organization (Molnár et al., 2008; Peng et al., 2024). Such experimental platforms may therefore provide a framework for systematically examining how stimulation parameters, cellular states, and pharmacological modulation interact to shape magnetic stimulation-induced plasticity in human cortical tissue.

Several limitations should be considered. The present study focused on excitatory synaptic plasticity in layer II/III pyramidal neurons and did not systematically examine inhibitory circuits, long-term network remodeling, or the molecular signaling pathways engaged by theta-burst magnetic stimulation in human tissue. Moreover, the experiments were performed in tissue obtained from a limited number of patients. Although the within-patient comparisons reduce the influence of interindividual variability and support a pulse-number-dependent differences in synaptic responses, larger and more diverse patient cohorts will be required to determine how factors such as disease state, cortical region, age, medication status, and intrinsic excitability shape responsiveness to magnetic stimulation across human cortical circuits (Huang (黄伟杰) et al., 2025; Lenz et al., 2024; Van Hoornweder et al., 2024). Future studies integrating electrophysiology with molecular, structural, and computational approaches may build on this experimental framework to further define how clinically relevant magnetic stimulation protocols engage plasticity mechanisms in the human brain.

## Materials and Methos

### Ethics statement

Human tissue was obtained from the local biobank of the Department for Neurosurgery at the Faculty of Medicine, University of Freiburg (AZ 472/15_160880) after informed consent was obtained from all individuals. All experiments were approved by the Local Ethics Committee, University of Freiburg (AZ 593/19). Patients gave their informed consent to use the neurosurgical resection material for research purposes.

### Preparation of acute human cortical slices

Human cortical tissue was obtained from resections performed for therapeutic purposes or to provide surgical access, as previously described (Straehle et al., 2023). None of the specimens exhibited macroscopically evident pathological alterations. Immediately after resection, the tissue was immersed in chilled, carbogenated NMDG-based artificial cerebrospinal fluid (NMDG-aCSF) composed of (in mM): 92 NMDG, 2.5 KCl, 1.25 NaH_2_PO_4_, 30 NaHCO_3_, 20 HEPES, 25 glucose, 2 thiourea, 5 sodium ascorbate, 3 sodium pyruvate, 0.5 CaCl_2_, and 10 MgSO_4_ (pH 7.3–7.4). The tissue was then embedded in low-melting-point agarose and sectioned at a thickness of 350 µm using a Leica VT1200S vibratome in chilled NMDG-aCSF. Following sectioning, slices were placed in 40-µm-pore cell strainers containing NMDG-aCSF at 34°C. The sodium concentration was subsequently increased in a stepwise manner (Ting et al., 2018). After recovery, slices were stored at room temperature until experimental use in extracellular solution containing (in mM): 92 NaCl, 2.5 KCl, 1.25 NaH_2_PO_4_, 30 NaHCO_3_, 20 HEPES, 25 glucose, 2 thiourea, 5 sodium ascorbate, 3 sodium pyruvate, 2 CaCl_2_, and 2 MgSO_4_ (pH 7.3–7.4). All solutions were continuously equilibrated with carbogen (95% O_2_/5% CO_2_).

**Table 1:**
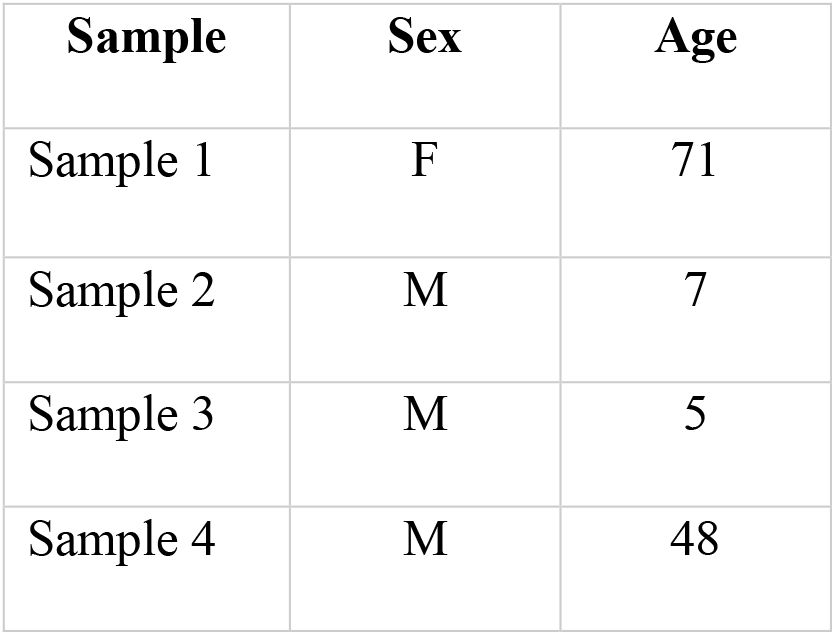
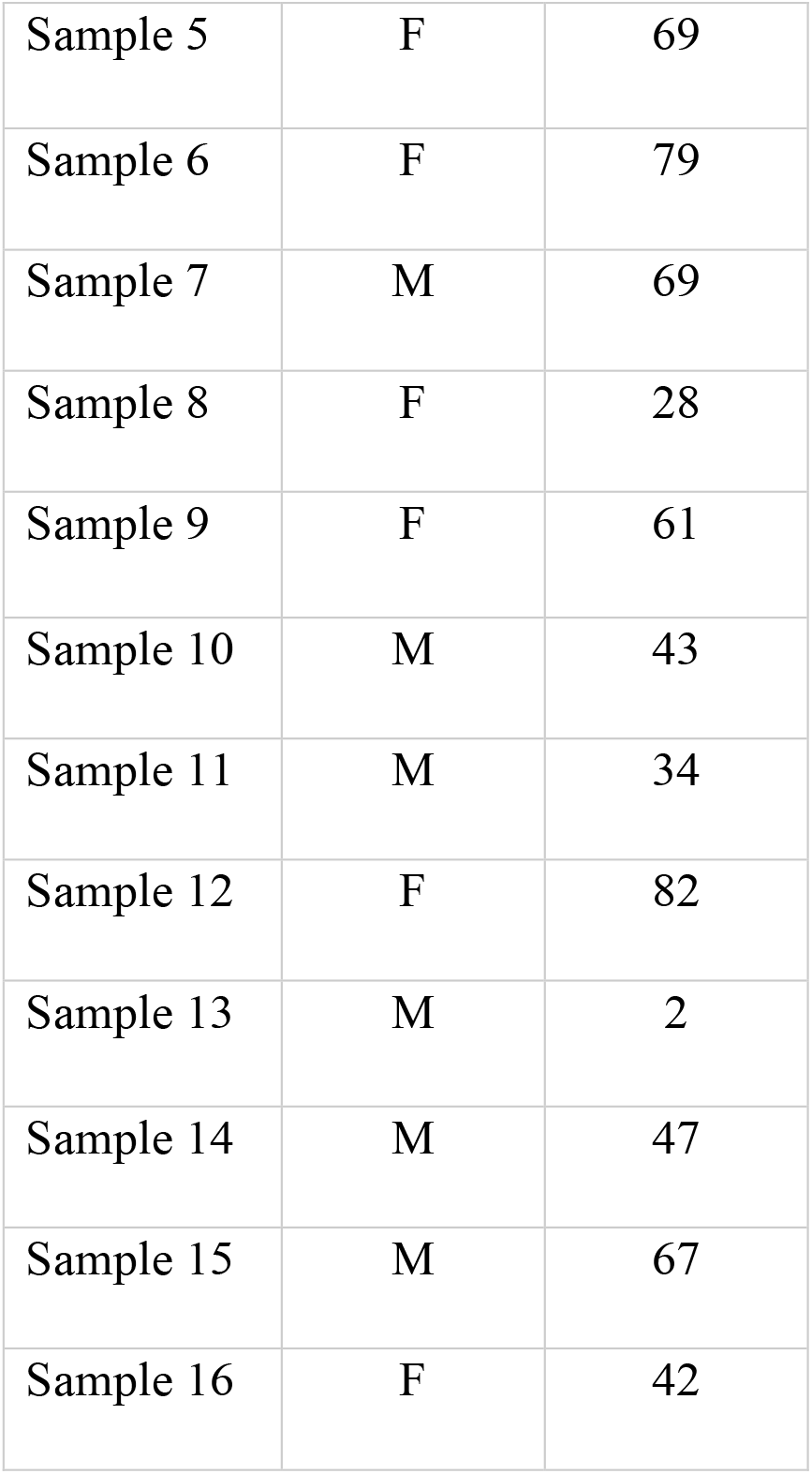
Human sample sex and age data.

| Sample | Sex | Age |
| --- | --- | --- |
| Sample 1 | F | 71 |
| Sample 2 | M | 7 |
| Sample 3 | M | 5 |
| Sample 4 | M | 48 |
| Sample 5 | F | 69 |
| Sample 6 | F | 79 |
| Sample 7 | M | 69 |
| Sample 8 | F | 28 |
| Sample 9 | F | 61 |
| Sample 10 | M | 43 |
| Sample 11 | M | 34 |
| Sample 12 | F | 82 |
| Sample 13 | M | 2 |
| Sample 14 | M | 47 |
| Sample 15 | M | 67 |
| Sample 16 | F | 42 |

### Electric field modeling

Macroscopic electric fields were computed using a three-dimensional finite element model with separate compartments for the bath solution and the tissue. The model geometry reproduced the dimensions of the in vitro setup, with the coil placed bellow the cultures at a distance of 1 mm from the Petri dish. Conductivity values were set to 1.654 S/m for the bath solution and 0.275 S/m for the tissue. The coil current rate of change was specified as 1.4 A/µs at 1% of maximum stimulator output (MSO) and scaled proportionally for higher stimulation intensities. Simulations were conducted in SimNIBS (version 3.2.6) and MATLAB (R2023a), using a validated model of a 70 mm MagStim figure-of-eight coil (Thielscher & Kammer, 2004). To obtain a robust estimate of the maximum electric field within the tissue, the 99th percentile of the field magnitude was calculated across the tissue volume.

### rMS ex vivo

Human neocortical slices were transferred to a standard 35-mm Petri dish containing extracellular solution composed of (in mM): 92 NaCl, 2.5 KCl, 1.25 NaH_2_PO_4_, 30 NaHCO_3_, 20 HEPES, 25 glucose, 2 thiourea, 5 sodium ascorbate, 3 sodium pyruvate, 2 CaCl_2_, and 2 MgSO_4_ (pH 7.3–7.4). The Petri dish was placed above a 70-mm figure-of-eight coil (D70 Air Film Coil, Magstim) connected to a Magstim Super Rapid^2^ Plus^1^ stimulator (Magstim), with the center of the dish aligned with the center of the coil. Slices received either an iTBS600 protocol (600 pulses; 20 cycles) or an iTBS900 protocol (900 pulses; 30 cycles). Each cycle comprised 10 bursts of three pulses delivered at 50 Hz, with bursts repeated at 5 Hz for 2 s, followed by an inter-train interval of 8 s, resulting in a total cycle duration of 10 s. Stimulation was applied at 50% of the maximum stimulator output (MSO), corresponding to an induced electric field strength of 90.5 V/m. All neocortical slices were oriented consistently, with the apical–basal axis of the cortical pyramidal neurons aligned with the direction of the induced electric field, thereby ensuring standardized stimulation geometry across experiments. Patient-matched sham slices underwent identical handling and experimental procedures but were positioned outside the effective magnetic field of the coil. Following stimulation, slices were returned to the continuously carbogenated storage solution and maintained at room temperature until further experimental assessment. The iTBS600 and iTBS900 experiments were performed as separate experimental series, each with its corresponding patient-matched sham control group.

### Electrophysiology

Whole-cell voltage-clamp recordings were obtained from visually identified pyramidal neurons in layers II/III of human neocortical slices at 35°C. Neurons were visualized using an Orca Fusion camera (Hamamatsu, Japan) mounted on an Olympus BX51 microscope (Olympus, Japan). During recording, slices were continuously superfused with carbogenated artificial cerebrospinal fluid containing (in mM): 126 NaCl, 2.5 KCl, 26 NaHCO_3_, 1.25 NaH_2_PO_4_, 2 CaCl_2_, 2 MgCl_2_, and 10 glucose. The solution was continuously equilibrated with 95% O_2_/5% CO_2_. To isolate miniature AMPA receptor-mediated excitatory postsynaptic currents (mEPSCs), 10 µM D-APV and 0.5 µM tetrodotoxin (TTX) were added to the bath solution. Patch pipettes were filled with an intracellular solution containing (in mM): 126 K-gluconate, 4 KCl, 4 Mg-ATP, 0.3 Na_2_-GTP, 10 phosphocreatine, and 10 HEPES, supplemented with 0.1% (w/v) biocytin. The pH was adjusted to 7.25 with KOH and the osmolarity to 290 mOsm with sucrose. Patch pipettes had resistances of 3–5 MΩ, and neurons were voltage-clamped at −70 mV. Liquid junction potentials were not corrected. Signals were acquired using a MultiClamp 700B amplifier and a Digidata 1550B digitizer (Molecular Devices), sampled at 10 kHz, and low-pass filtered at 6 kHz using a Bessel filter. Series resistance was assessed at 2-min intervals throughout each recording, and recordings were discontinued if it reached or exceeded 30 MΩ. Leak current was monitored continuously; no recordings showed changes requiring exclusion. Recordings from patient-matched sham- and iTBS-treated slices were performed 2–4 h after stimulation in a pseudorandomized order to minimize acquisition bias.

### Fluorescent labeling of recorded neurons

Following whole-cell recordings, human neocortical slices containing biocytin-filled pyramidal neurons were fixed for 1 h in 0.01 M PBS containing 4% (w/v) paraformaldehyde and 4% (w/v) sucrose. After washing with 0.01 M PBS, the slices were incubated for 4 h with Alexa Fluor 488-conjugated streptavidin (1:1000) diluted in 0.01 M PBS containing 10% (v/v) normal goat serum and 0.1% (v/v) Triton X-100. The slices were subsequently washed with 0.01 M PBS, transferred onto glass slides, and coverslipped using an antifade mounting medium. Confocal images were acquired using a Leica TCS SP8 laser-scanning microscope equipped with 20× (NA 0.75), 40× (NA 1.30), and 63× (NA 1.40) oil-immersion objectives.

### Experimental design and statistical analysis

Analyses were performed with the person analyzing the data blind to the experimental condition. Whole-cell patch clamp data were analyzed using pClamp 11.2 software suite (Molecular Devices) and Easy Electrophysiology 2.5.0.2 (Easy Electrophysiology Ltd.). Statistical comparisons were performed using Mann–Whitney tests and Kruskal-Wallis tests (GraphPad Prism 7). All values reported in the figures and main text are expressed as mean ± SEM. p <0.05 was considered statistically significant.

### Digital illustrations

Confocal image stacks were exported as 2D projections and stored as TIFF files. Figures were prepared using the Affinity software and Biorender.

## Acknowledgments

The authors thank Lejla Kardic and Brunhilde Baumer for their valuable technical assistance with solution preparation and the transfer of human cortical tissue from the operating room to the laboratory.

